## Supplementary material for "The role of ventromedial prefrontal cortex in reward valuation and future thinking during intertemporal choice": Table 1

Table 1. vmPFC patients’ demographic and clinical data.

| vmPFC patient | Age  (y) | Edu  (y) | Sex  (y) | Time since lesion  (y) | EFT Int  (z score) | EFT Ext  (z score) | PF | LF | DS | LL  Imm | LL  Del | ROCF Copy | ROCF  Recall |
| --- | --- | --- | --- | --- | --- | --- | --- | --- | --- | --- | --- | --- | --- |
| P1 (I) | 55 | 13 | M | 4 | -1.42 | 0.58 | 21% | 23% | 34% | 14% | 17% | 100% | 50% |
| P2 (I) | 46 | 13 | M | 7 | -1.54 | -1.44 | 38% | 7% | 49% | 12% | 8% | 100% | 41% |
| P3 (I) | 56 | 8 | M | 13 | -1.43 | -0.73 | 42% | 16% | 23% | 0.43% | 3% | 25% | 2% |
| P4 (I) | 57 | 8 | M | 7 | -1.57 | -0.28 | 42% | 31% | 23% | 7% | 12% | 89% | 27% |
| P5 (C) | 58 | 15 | F | 8 | - | - | 82% | 35% | 18% | 1% | 0.02% | 2% | 13% |
| P6 (C) | 76 | 16 | F | 5 | 0.20 | -0.86 | 55% | 40% | 80% | 81% | 50% | 67% | 62% |
| P7 (C) | 54 | 13 | F | 2 | -2.27 | -1.64 | 58% | 30% | 59% | 2% | 2-3% | 8% | 42% |
| P8 (C) | 65 | 18 | M | 4 | -1.93 | -1.00 | 45% | 2% | 39% | 8% | 7% | 22% | 18% |
| P9 (C) | 56 | 20 | M | 4 | -2.24 | -1.24 | 47% | - | 39% | 1% | 0.7% | 68% | 1% |
| P10 (C) | 51 | 10 | M | 8 | - | - | 45% | 20% | 59% | 4% | 0.7% | 84% | 13% |
| P11 (C) | 66 | 15 | F | 1 | -1.73 | -1.41 | 47% | 55% | 39% | 1% | 1% | 70% | 2% |
| P12 (C) | 49 | 12 | M | 5 | -1.89 | 0.95 | 86% | 50% | 39% | 1% | 0.03% | 58% | 0.7% |

Note. (I) = patient tested in Italy; (C) = patient tested in Canada; M = male; F = female; Edu = education; y = years; vmPFC = ventromedial prefrontal cortex; EFT Int = internal details at the Crovitz episodic future thinking task; EFT Ext = external details at the Crovitz episodic future thinking task; PF = Premorbid functioning, **based on the Full scale IQ at Wechsler Abbreviated Scale of Intelligence (WAIS–IV; Wechsler, 2008) for Canadian patients P7, P9, P12, on the Wechsler test of adult reading (WTAR; Holdnack, 2001) for Canadian patients P6 and P11, on the National Adult Reading Test (NART) (Paolo and Ryan, 1992) for Canadian patients P5, P8, and P10,** and on the Raven Standard Progressive Matrices (SPM) for all Italian patients (Spinnler and Tognoni, 1987); LF = Letter fluency (Spinnler and Tognoni, 1987; Spreen and Strauss, 1998); DS = Digit span **(forward)**; LL Imm = List learning immediate recall, LL Del = List learning delayed recall, assessed with the Buschke–Fuld Test (Buschke and Fuld, 1974; Spinnler and Tognoni, 1987) in Italian patients, and with the California Verbal Learning Test-II (Woods et al., 2006) in Canadian vmPFC patients; ROCF = Rey-Osterrieth Complex Figure (Spinnler and Tognoni, 1987; Spreen and Strauss, 1998). For PF, LF, DS, LL, and ROCF we report percentile scores. Dashes indicate missing data.
